# Intrinsic plasticity of cerebellar Purkinje cells in motor learning circuits

**DOI:** 10.1101/513283

**Authors:** Dong Cheol Jang, Hyun Geun Shim, Sang Jeong Kim

## Abstract

Intrinsic plasticity of cerebellar Purkinje cells (PCs) is recently highlighted in the cerebellar local circuits, however, its physiological impact on the cerebellar learning and memory remains elusive. Using a mouse model of memory consolidation deficiency, we found that the intrinsic plasticity of PCs may be involved in motor memory consolidation. Gain-up training of the vestibulo-ocular reflex produced a decrease in the synaptic weight of PCs in both the wild-type and knockout groups. However, intrinsic plasticity was impaired only in the knockout mice. Furthermore, the observed defects in the intrinsic plasticity of PCs led to the formation of improper neural plasticity in the vestibular nucleus (VN) neurons. Our results suggest that the synergistic modulation of intrinsic and synaptic plasticity in PCs is required for the changes in following plasticity in the VN, thereby contributes to the long-term storage of motor memory.

## Introduction

It is widely believed that the cellular basis of memory is derived from modifications of synaptic transmission, such as long-term potentiation (LTP) and long-term depression (LTD) (Kandel, Dudai, & Mayford, 2014). For decades, since Ito proposed the flocculus hypothesis (Ito, 1982), numerous studies have demonstrated that synaptic plasticity between parallel fibers (PFs) and cerebellar Purkinje cells (PCs) is the key mechanism of vestibule-ocular reflex (VOR) adaptation (De Zeeuw et al., 1998; Hansel et al., 2006; Inoshita & Hirano, 2018; Kakegawa et al., 2018; Schonewille et al., 2010). However, several works proposed that the synaptic plasticity at the PF-PC synapse is not sufficient to explain motor learning (Ito, 2013; Ke, Guo, & Raymond, 2009; Schonewille et al., 2011; Wulff et al., 2009). Emerging evidence suggests that neural plasticity at multiple sites, including the cerebellar cortex and vestibular nucleus (VN), is required for motor learning (Boyden, Katoh, & Raymond, 2004; Clopath, Badura, Zeeuw, & Brunel, 2014; Porrill & Dean, 2007; Yamazaki, Nagao, Lennon, & Tanaka, 2015). Furthermore, it has been suggested that motor memory is formed in cortical areas through PF-PC plasticity at the early phase of adaption and transferred to sub-cortical area, especially in the VN (Ito, 2013). In the motor learning circuits, cerebellar PCs integrate the information and then project their output signal to the VN. Interestingly, synaptic plasticity at mossy fibers (MFs) to VN neurons show dependency on the activity of PCs (McElvain, Bagnall, Sakatos, & Lac, 2010; Medina, 2010). This suggests that the activity-dependent modulation of the output of PCs might derive adequate memory transduction from the cerebellar cortex to the VN. Indeed, the changes in intrinsic neuronal excitability (intrinsic plasticity) has been suggested as additional mechanism for memory storage (Daoudal & Debanne, 2003; Zhang & Linden, 2003). However, the impact of intrinsic plasticity on the information storage is still elusive. we previously verified that the excitability of PCs shows bi-directionality in response to the different patterns of synaptic plasticity induction (Shim et al., 2017) and postulated that the intrinsic plasticity of PCs plays a role in VOR adaptation, based on former results (Ryu et al., 2017).

Here, we provide insight into the circuit mechanism through which the intrinsic plasticity of cerebellar PCs may be required for long-term memory storage. We adopted cerebellum-dependent eye movement learning paradigm, and recorded both synaptic and intrinsic plasticity in the cerebellar PCs and the VN neurons. By dissecting the time after training, we found both type of plasticity had been induced by the training in both regions. Furthermore, each plasticity showed different temporal dynamics through the period. Simultaneously, we recorded the same plasticity in memory consolidation deficit mouse model, in which stromal interaction molecule 1 (STIM1) was knocked out PC specifically (STIM1^PKO^) (Ryu et al., 2017), to investigate the relationship between the plasticity and consolidation process. Interestingly, the STIM1^PKO^ mice showed deficient intrinsic plasticity, although synaptic plasticity was induced after learning. Furthermore, neither synaptic plasticity nor intrinsic plasticity of the VN neurons was observed in STIM^PKO^ mice after training. This implies that the subsequent increases in the neural activity in the VN neurons may be derived from changes in the cortical output, as determined by the intrinsic plasticity of the cerebellar PCs.

## Results

### Consolidation deficit in the STIM1^PKO^ mouse may occur between 1 to 4 hours after learning

Accumulating evidence supports the theory that eye movement memory is firstly formed in the cerebellar cortex and is then transferred to the brainstem (Ito, 2013; Kassardjian et al., 2005; Matsuno et al., 2016; Okamoto, Endo, Shirao, & Nagao, 2011a). This sequential memory acquisition within the cortex and brainstem has been implicated in long-term memory storage. Interestingly, Okamoto et al., (2011a) observed that memory transfer occurs between 2.5 and 4 hours after learning in mice. This implies that this period of time is the critical period for communication between the cerebellar cortex and sub-cortical areas (in this circumstance, the VN). Because we have previously reported that an impairment in long-term memory storage, but not memory acquisition, is observed in the STIM1^PKO^ mice (Ryu et al., 2017), we expected that a dysfunctional memory transfer process would manifest as a consolidation deficit in this model. To determine whether VOR memory was attenuated in the STIM1^PKO^ mice over the memory transfer period, we selected three time periods after learning task in which measurements were made: 0.5 and 1 hours two points as short-term period, 4 hours as the mid-term period and 24 hours as the long-term period. The memory retention level was evaluated at each of these periods. Both groups showed normal basal oculomotor performance (Figure 1-figure supplement 1) and no memory impairment at the short-term period (Figure 1B). However, at the mid-term period, the memory retention level was significantly lower in the STIM1^PKO^ group than in the wild-type group. Furthermore, at the long-term period memory retention was eliminated in the STIM1^PKO^ group (Figure 1B). This temporal alteration in the memory retention level was re-calculated as the ratio of the remaining memory at the test session to the acquired memory at the training session (Figure 1C). This revealed a showing gradual reduction in memory retention over the studied periods. Given that the slight decline in the level of memory retention began 1 hour after the learning task and developed further, we speculated that the impaired motor memory consolidation that was observed in the STIM1^PKO^ mice was based on defect of memory transfer process, leading to inappropriate communication between the cerebellar cortex and the VN.

**Figure 1.**
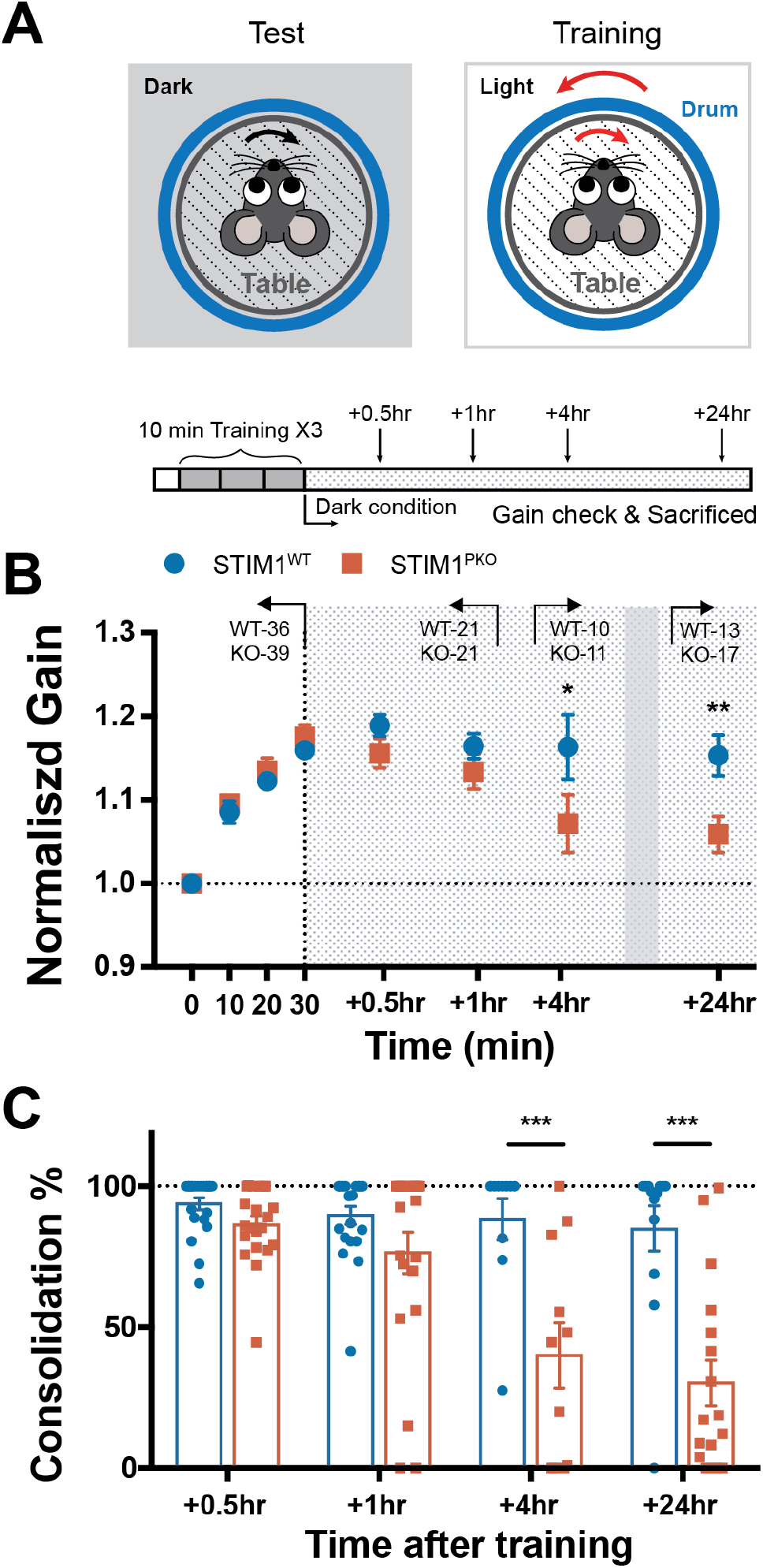
Long-term memory storage was impaired in STIM1^PKO^ mice. (A) Illustration of the VOR test and training (upper) and experimental scheme (below). For training, visuo-vestibular stimulation was delivered through drum and turntable, 10 min each session, three times; dVOR at 0.5 Hz was recorded before the learning and after each session of training. After whole training session, animals were put in the completely dark condition. (B)Normalized gain of the eye movement in learning. Note there is no significant differences between wild-type and STIM1^PKO^ mice in learning (points on white-colored background; wild-type n=36, STIM1^PKO^ n=39). We measured memory retention level at 0.5 and 1 hour (described as short-term; wild-type n=21, STIM1^PKO^ n=21), 4 hours (mid-term; wild-type n=10, STIM1^PKO^ n=11) and 24 hours (long-term period; wild-type n=13, STIM1^PKO^ n=17) after training (points on grey-dotted background). STIM1^PKO^ showed significantly lower memory retention level from the mid-term period compared to the wild-type littermates (4hr, p=0.037; 24hr; p=0.004). (C) Calculated consolidation level. Memory retention level was obtained by calculating the ratio remained to learned memory (4hr, p <0.001; 24hr, p < 0.001). Two-way repeated measure ANOVA was used for panel B, and asterisks in the graph were marked by post-hoc Sidak test for pairwise comparison. Unpaired *t*-test was used for panel C. Error bars denote SEM, *p < 0.05, **p < 0.01, ***p < 0.001.

### Learning generally induces PF-PC synaptic plasticity in both groups

Many previous studies, using various LTD-deficient animal models, have reported that synaptic plasticity at the PF-PC synapse is strongly correlated with motor learning (Boyden et al., 2006; De Zeeuw et al., 1998; Hansel et al., 2006). Optokinetic response (OKR)-induced synaptic LTD in the cerebellar cortex has been recently reported (Inoshita & Hirano, 2018), but plasticity induced by the VOR adaptation has yet to be proved. Late-phase LTD has also been implicated in VOR memory consolidation (Ahn, Ginty, & Linden, 1999; Boyden et al., 2006). This suggests that a learning-induced long-lasting reduction of cerebellar cortical activity drives the transduction of memory to the sub-cortical region. Interestingly, because the time course of late-phase LTD is similar to that of the mid-term period of study, we investigated whether PF-PC LTD is involved in memory consolidation. To verify this in detail, electrophysiological *ex vivo* recordings were made from floccular PCs to investigate the signs of synaptic plasticity at the short-, mid-and long-term periods after learning task. This approach enabled us to monitor neuronal activity for periods of over an hour, overcoming the experimental limitation of the whole-cell patch clamp technique. Due to the location of micro zone of the flocculus that regulates horizontal VOR behavior, we recorded from PCs that were located in the medial part of the flocculus (Schonewille et al., 2006). As a control group, we used sham animals that had undergone surgery and restraint without the learning task. We firstly measured the PF-stimuli-evoked synaptic response (eEPSC) following the injection of ranges of electrical stimuli intensities. In these experiments, the amplitude of the eEPSC was decreased at the short-(1hr) and mid-term (4hr) period than sham group. However, this alteration was recovered, with the eEPSC returning to a level that was not significantly different to that of sham control before the long-term phase in both group (Figure 2B and C). In addition to the PF-evoked synaptic events, the spontaneous excitatory postsynaptic current (sEPSC) was recorded after the learning task. Between the sham groups of both genotypes, there was no significant difference in the spontaneous glutamatergic synaptic transmission (Figure 2-figure supplement 1). The distribution of the inter-event intervals (IEIs) of synaptic events was shifted to the right after the learning task and was restored at the long-term period in both groups (Figure 2E and F). However, the mean frequency of sEPSC in the STIM1^PKO^ mice was transiently reduced until 1 hour after the learning task, and was recovered 4 hours later, while in the change of wild-type littermates, the change was maintained throughout all of the time periods after the learning task (insets of Figure 2E and F). Conversely, the aspect of changes in the sEPSC amplitudes in the STIM1^PKO^ mice were comparable to that of the wild-type littermates (Figure 2F and I). In both groups, the distribution of the sEPSC amplitude was left-shifted after the learning task and this change was maintained until the long-term period. Collectively, these results, as well as the results of a previous study (Boyden et al., 2006), indicate that VOR gain-up training elicited synaptic weakening of the PF-PC synapses. Although the alterations of sEPSC frequency that were observed in the STIM1^PKO^ mice were relatively transient compared to the results observed in the wild-type group, overall, the aspects of plasticity that were induced by VOR adaptation were similar between the groups. Therefore, we suggest that the learning-induced PF-PC LTD presumably did not contribute to the long-term memory deficit observed in the STIM1^PKO^ mice.

**Figure 2.**
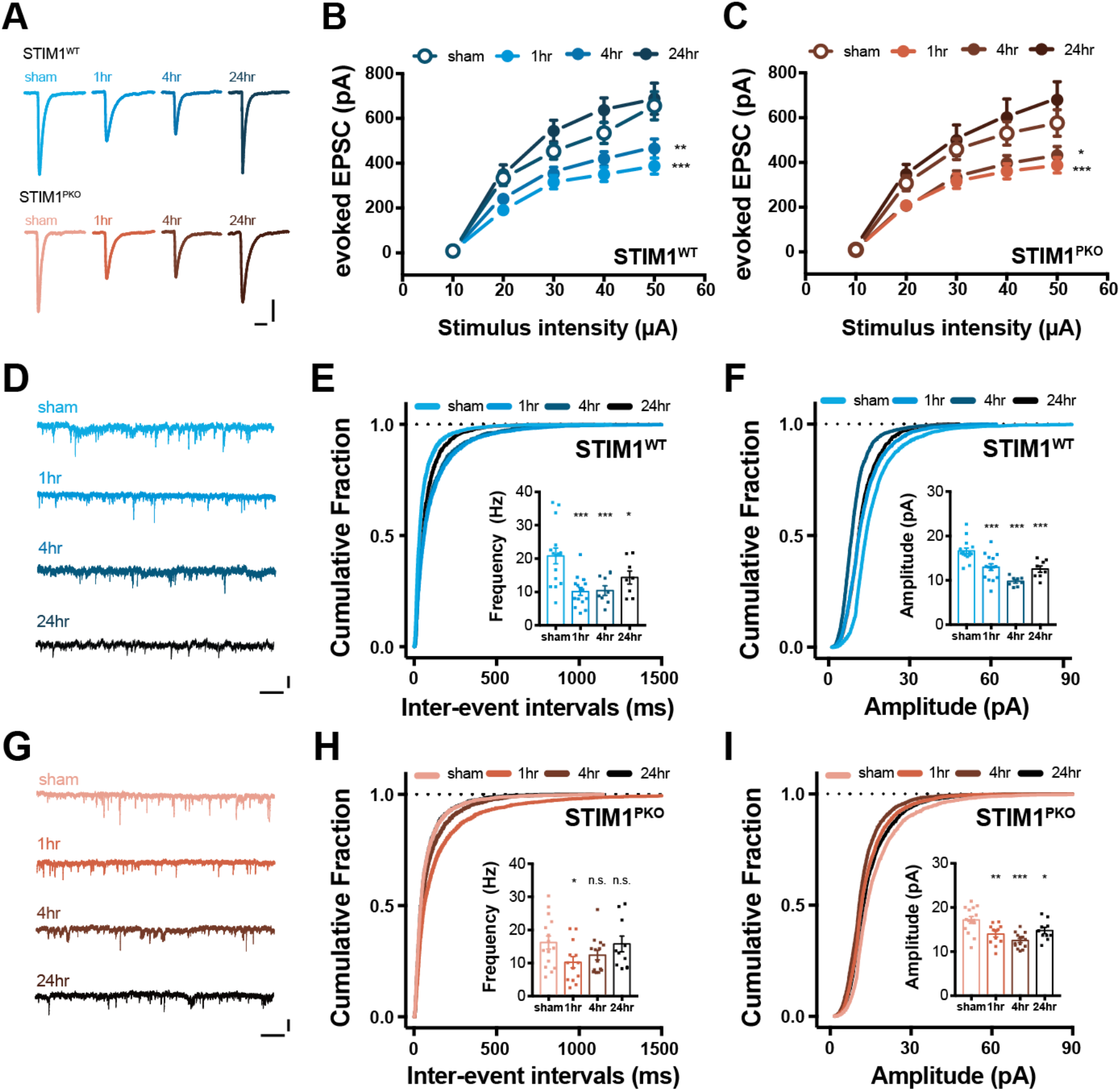
Long-term depression at the PF-PC synapses after gain-up learning. (A) Representative eEPSCs traces of both genotype groups in each time point. Scale bars, 200 pA (vertical) and 30 ms (horizontal). (B) Amplitude of eEPSC by serial PF stimulation in wild-type littermates. In comparison to sham group (n=9), the amplitude was significantly reduced at 1 hour and 4 hours after learning (at 50 µA injection; 1hr, n=19, p < 0.001; 4hr, n=9, p=0.002), and depressed amplitude was recovered 24 hours after (at 50 µA injection; 24hr, n=8, p=0.922). (C) Amplitude of eEPSC by serial PF stimulation in STIM1^PKO^. Same as wild-type littermates, the amplitude was considerably decreased at 1 and 4 hours after, and restored at 24 hours after learning (at 50 µA injection; sham, n=14; 1hr, n=14, p < 0.001; 4hr, n=14, p=0.013; 24hr, n=7, p=0.232). (D) Representative sEPSC traces of wild-type group in each time point. Scale bars, 25 pA (vertical) and 1s (horizontal). (E) Cumulative plots of inter-events-interval (IEI) of sEPSC in wild-type littermates. The cumulative fraction was right-shifted after learning, implying reduction of frequency (sham n=15, 1hr n=15, 4hr n=8, 24hr n=8). Inset bar graph is mean frequencies of sEPSC indicating depression of frequency was maintained until 24 hours after learning in comparison to sham group (1hr, p < 0.001; 4hr, p < 0.001; 24hr, p=0.029). (F) Cumulative plots of amplitude of sEPSC in wild-type littermates. The cumulative fraction was left-shifted after learning, implying reduction of amplitude. Inset bar graph is mean amplitudes of sEPSC indicating that depression of amplitude was maintained for 24 hours after learning (1hr, p <0.001; 4hr, p < 0.001; 24hr, p < 0.001). (G) Representative sEPSC traces of STIM^PKO^ group in each time point. Scale bars, 25 pA (vertical) and 1s (horizontal). (H) Cumulative plots of inter-events-interval (IEI) of sEPSC in STIM^PKO^ mice. The cumulative fraction was right-shifted 1 hour after learning, but most of changes returned to sham level from 4 hours after training (sham n=15, 1hr n=13, 4hr n=14, 24hr n=10). Inset shows summarizing bar graph of sEPSC frequency, implying that frequency of sEPSC was transiently depressed at 1 hour and recovered from 4 hours after learning (1hr, p <0.001; 4hr, p < 0.001; 24hr, p < 0.001). (I) Cumulative plots of amplitude of sEPSC in STIM^PKO^ mice. The cumulative fraction was left-shifted after learning. Inset bar graph shows that amplitude of sEPSC was depressed and maintained for 24 hours after learning (1hr, p=0.001; 4hr, p < 0.001; 24hr, p=0.016). Two-way repeated measure ANOVA was used for panel A and B, and asterisks in the graph were marked by post-hoc Sidak test for pairwise comparison. One-way ANOVA with Fisher LSD test was used for insets in panel C, D, E and F. Asterisks in each time points were calculated by comparing to sham groups. Error bars denote SEM. *p < 0.05, **p < 0.01, ***p < 0.001.

### Learning-induced intrinsic plasticity shows relevance in memory consolidation

Experience-dependent neural plasticity includes not only synaptic plasticity but also alterations in the intrinsic excitability (Daoudal & Debanne, 2003; Zhang & Linden, 2003). Because PF-PC LTD was not sufficient to account memory consolidation deficit in STIM1^PKO^, we investigated whether if intrinsic plasticity, the other form of neural plasticity, could be a considerable factor in memory consolidation. In the light of our previous results (Ryu et al., 2017), excitability of PC is highly suspicious, because basal excitability of PCs was significantly reduced in STIM1^PKO^ but there were no developmental differences, such as the morphology of PCs, expression level of Ca^2+^-related channels and CF-induced complex spike. To investigate whether excitability changes of cerebellar PCs are required for memory consolidation, we firstly performed whole-cell patch clamp recordings to compare the long-term depression of intrinsic excitability (LTD-IE) in the STIM1^PKO^ group and the wild-type littermates. A PF burst protocol (7 of 100 Hz PF burst followed by a single CF stimulation; Shim et al., 2017) was introduced to induce PC synaptic and intrinsic plasticity in the presence of an inhibitory synaptic transmission inhibitor, picrotoxin. As shown in Figure 2, both STIM1^PKO^ and the wild-type groups showed normal induction of PF-PC LTD (Figure 3-figure supplement 1A) and LTD-IE (Figure 3-figure supplement 1B). However, the magnitude of the intrinsic plasticity was weaker in the STIM1^PKO^ group than in the wild-type group. Interestingly, the reduction in the excitability following LTD induction was recovered 40 min after the induction in the STIM1^PKO^ mice, whereas, in the wild-type littermates LTD-IE was elicited and developed further (Figure 3-figure supplement 1B).

Next, we examined the temporal alteration of PC excitability through *ex vivo* recordings after the learning task; at short-, mid-and long-term time periods. In agreement with the results of *in vitro* experiments (Shim et al., 2017; Figure 3-figure supplement 1B), the firing frequency was decreased 1 hour after training in the wild-type littermates (Figure 3A and B). The AP firing frequency of PCs was measured in current clamp mode through the injection of brief current steps from the membrane potential of approximately −70 mV (500 ms, from +100 pA to + 500 pA with an increment of 100 pA, step interval 4.5 s). The learning-induced intrinsic plasticity was partially recovered at the mid-term time period and fully recovered to the value from the sham control at the long-term period. However, the STIM1^PKO^ group showed a deficiency in the learning-induced intrinsic plasticity throughout the studied periods (Figure 3C and D). Comparing the results from the different genotypes over the same time period, wild-type littermates has significantly higher firing frequency in sham control as we reported (Ryu et al., 2017), but the frequency reversed in the short-term period and gradually recovered over the time period studied (Figure 3-figure supplement 1C-F). Given the notion that the intrinsic plasticity of cerebellar PCs amplifies the modification of synaptic weight to properly project the learned signal from the PCs to their relay neurons (Shim et al., 2017), our observation may imply that the impairment of intrinsic plasticity results in dysfunctional memory transfer to the VN neurons. Because intrinsic plasticity was almost restored 40 min after plasticity induction in the STIM1^PKO^ group (Figure 3-figure supplement 1B), we expect that the learning-induced intrinsic plasticity may be already abolished within 1 hour after training. Taken together, we conclude that the intrinsic plasticity of the cerebellar PCs would be involved in long-lasting reduction of the cerebellar cortical activity, and thereby, may contribute to VOR memory consolidation.

**Figure 3.**
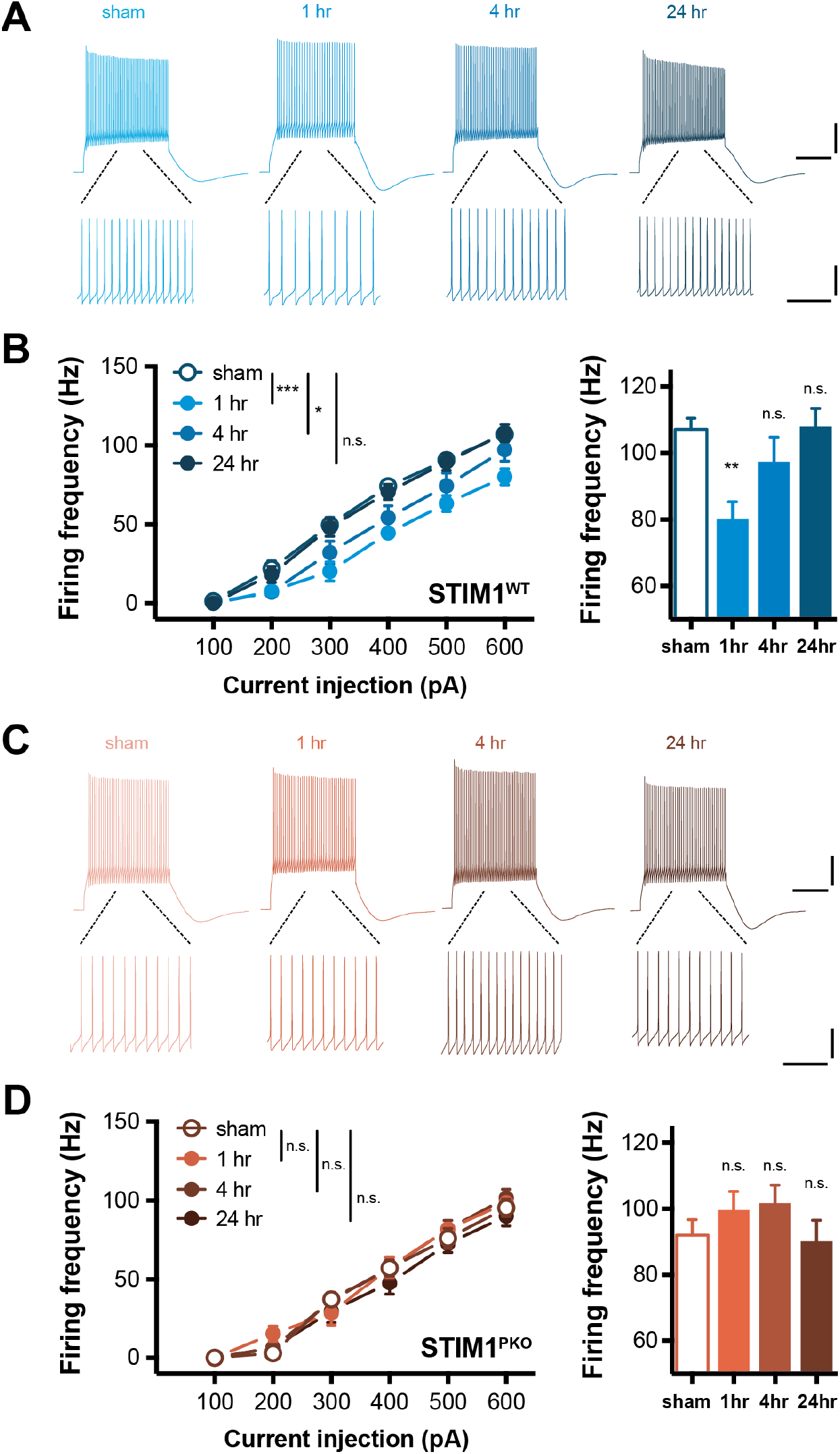
VOR adaptation also induces plastic change of intrinsic excitability in PCs. (A) Representative traces from whole-cell recording in wild-type group. Scale bars (upper), 20 mV (vertical) and 200 ms (horizontal). Scale bars (lower), 20 mV (vertical) and 50 ms (horizontal). (B) PC excitability of wild-type littermates over several time points. VOR adaptation decreased gain responses of the cerebellar PCs in response to square-wised current injection ranging from 100 pA to 600 pA for 500 ms (sham vs 1hr, p < 0.001; sham vs 4hr, p=0.024; sham vs 24hr, p=0.717, left; sham, n=20; 1hr, n=11; 4hr, n=16; 24hr, n=20). Excitability in 600 pA injection was significantly decreased at short-term (1hr, p=0.002) and mostly recovered at mid-term (4hr, p=0.196) and fully recovered at long-term (24hr, p=0.914). (C) Representative traces from whole-cell recording in STIM1^PKO^ group. Scale bars (upper), 20 mV (vertical) and 200 ms (horizontal). Scale bars (lower), 20 mV (vertical) and 50 ms (horizontal). (D) Excitability of PC in STIM1^PKO^. Different from wild-type littermates, STIM1^PKO^ showed no alteration of excitability after learning (sham vs 1hr, p=370; sham vs 4hr, p=0.343; sham vs 24hr, p=0.768, left; sham, n=17; 1hr, n=13; 4hr, n=17; 24hr, n=13). There were no significant changes in 600 pA injection (1hr, p=0.371.; 4hr, p=0.184; 24hr, p=0.814) Two-way repeated measure ANOVA was used for injected current-frequency graphs. One-way ANOVA with Fisher LSD test was used for bar graphs. Asterisks in each time points were calculated by comparing to sham groups. Error bars denote SEM. **p < 0.01.

### Appropriate synaptic and intrinsic plasticity of VN neurons require intrinsic plasticity of cerebellar PC

We observed an impairment of intrinsic plasticity in the cerebellar PCs of the STIM1^PKO^ group through *in vitro* and *ex vivo* recordings. This suggests that memory consolidation requires the transduction of a memory from the cerebellar cortex into the brainstem through the intrinsic plasticity of PCs. A large population of VN neurons receive information from floccular PCs (Matsuno et al., 2016; Shin et al., 2011). Furthermore, the output of the cerebellar PCs serve as an instructive signal in the neuronal plasticity between MFs and VN neurons (Clopath et al., 2014; Dean, Porrill, Ekerot, & Jörntell, 2010; McElvain et al., 2010; Medina, 2010; Porrill & Dean, 2007; Shutoh, Ohki, Kitazawa, Itohara, & Nagao, 2006; Yamazaki et al., 2015). Thus, we hypothesized that the impairment of the intrinsic plasticity of the cerebellar cortex that was observed in STIM1^PKO^ mice would lead to an inadequate alteration of the VN neuron activity following VOR adaptation. To address this, we performed *ex vivo* recordings to investigate spontaneous synaptic transmission after the learning task during three distinct time periods, short-, mid- and long-term by *ex vivo* recordings. In the sham groups, the frequency of sEPSC in the STIM1^PKO^ group showed remarkable augmentation compared to the wild-type group (Figure 4-figure supplement 1A). However, the sEPSC amplitude was not significantly different between the STIM1^PKO^ group and wild-type group (Figure 4-figure supplement 1B). These results imply that the homeostatic scaling in the VN neurons are due to the reduction of PC excitability in the STIM1^PKO^ group (Figure 3-figure supplement 1C). Intriguingly, synaptic transmission was found to be potentiated after VOR adaptation in the wild-type littermates throughout the periods of study (Figure 4A-C). Although the increase in the mean frequency of sEPSC after training seemed to be restored at the long-term period, the cumulative distribution of the IEIs was found to be left-shifted throughout the periods of study (Figure 4B). The cumulative fraction of the sEPSC amplitude was especially right shifted at the long-term period, indicating that the proportion of increased glutamatergic synaptic events was enhanced during this period (Figure 4C). However, the mean value was not significantly altered compared to that of the sham control (Figure 4C, inset). These results indicate that VOR gain-up learning induces LTP at the MF- VN synapse, in line with the previous expectation (Boyden et al., 2006). In contrast to the results presented from the wild-type littermates, the STIM1^PKO^ group showed a slight depression of sEPSC frequency in cumulative distribution in the short- and mid-term time periods that continuously recovered to baseline (Figure 4E). However, the mean frequency was not significantly altered among the periods of study (Figure 4E, inset). The amplitude of sEPSC was slightly left-shifted in the short- and mid-term time periods, and mean amplitude in mid-term time period was significantly lower than that of the sham group (Figure 4F). In light of previous reports, which have suggested that cerebellar PC activity contribute to MF-VN plasticity (Dean et al., 2010; Matsuno et al., 2016; McElvain et al., 2010; Medina, 2010), we speculated that the synaptic plasticity at the MF-VN synapse is inappropriately induced in the STIM1^PKO^ group due to the absence of PC intrinsic plasticity.

**Figure 4.**
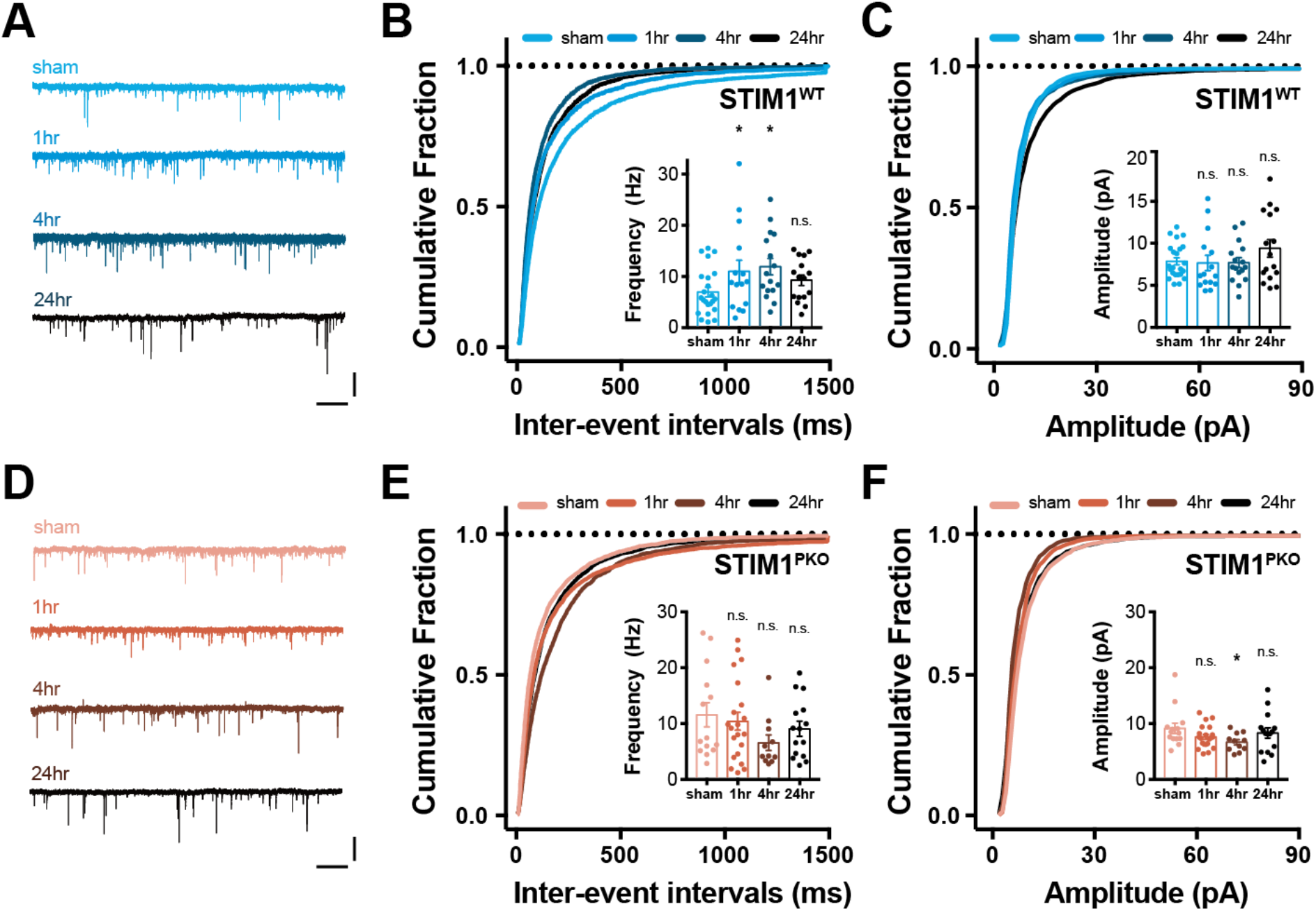
VOR gain-up learning induced long-term potentiation of the excitatory input in VN neurons. (A) Representative sEPSC traces of wild-type group in each time point. Scale bars, 25pA (vertical) and 1s (horizontal). (B) Inter-events-interval (IEI) of sEPSC in wild-type mice. The cumulative distributions of IEI were left-shifted after learning (sham, n=23; 1hr, n=15; 4hr, n=16; 24hr, n=16). The frequency of sEPSC was potentiated at short-(1hr) and mid-term (4hr) after training (inset bar graph; 1hr, p=0.046; 4hr, p=0.013; 24hr, p=0.242). (C) Amplitude of sEPSC in wild-type mice. The cumulative distribution was right-shifted at 24 hours after learning. There was trend of potentiation at long-term (24hr) after training, but overall, the amplitude of sEPSC was not significantly affected by learning (inset bar graph; 1hr, p=0.850; 4hr, p=0.874; 24hr, p=0.122). (D) Representative sEPSC traces of STIM1^PKO^ group in each time point. Scale bars, 25 pA (vertical) and 1s (horizontal). (E) Inter-events-interval (IEI) of sEPSC in STIM^PKO^ mice. In contrast to wild-type littermates, the cumulative distribution was slightly right-shifted after learning (sham, n=14; 1hr, n=21; 4hr, n=111; 24hr, n=15). The mean frequency of sEPSC was not significantly altered after learning (inset bar graph; 1hr, p=0.613; 4hr, p=0.066; 24hr, p=0.314). (F) Cumulative plots of amplitude of sEPSC in STIM^PKO^ mice. The cumulative distribution was slightly left-shifted at 4 hours after learning. The amplitude of sEPSC was significantly reduced at mid-term (4hr) after learning (inset bar graph; 1hr, p=0.111; 4hr, p=0.030; 24hr, p=0.420). One-way ANOVA with Fisher LSD test was used for bar graphs in all panels. Asterisks in each time points were calculated by comparing to sham groups. Error bars denote SEM. *p < 0.01

Furthermore, we asked whether the intrinsic plasticity of cerebellar PCs is also required for the adequate induction of intrinsic plasticity in the VN neurons, because VOR training involves a change in the excitability as well as synaptic transmission (Carcaud et al., 2017; Shutoh et al., 2006). To answer this, the gain responses were measured through the injection of square-wised somatic depolarizing current into the VN neurons at the three time periods after the learning task. The VN neurons of the STIM1^PKO^ group showed higher firing frequency in response to the current injection than were observed in the wild-type littermates in the sham group (Figure 4-figure supplement 1C). Interestingly, VOR training elicited the intrinsic plasticity of the VN neurons in the wild-type littermates (Figure 5A and B), whereas, there was no alteration of the gain responses in the STIM1^PKO^ group (Figure 5C and D). The excitability of the VN neurons gradually increased over the studied time periods and the intrinsic plasticity was maintained 24 hours after training (Figure 5A). The difference between the sham groups of both genotypes was faded during the short- and the mid-term time period, and finally reversed with statistical significance at the long-term time period (Figure 4-figure supplement 1C-F). To clarify whether the neural plasticity in the VN neurons is affected by knockout of STIM1 in the PC, we delivered the conventional protocol for induction of LTP in VN neurons. The VN neurons from the wild-type and STIM1^PKO^ groups exerted potentiation of the synaptic weight and excitability (Figure 5-figure supplement 1). Taken together with modification of synaptic weight and intrinsic properties in VN neurons, we suggest that the intrinsic plasticity of the cerebellar PCs following VOR adaptation could enable to induction of the proper forms of neuronal plasticity in VN neurons, corresponding to the specific behavior, such as consolidation of memory.

**Figure 5.**
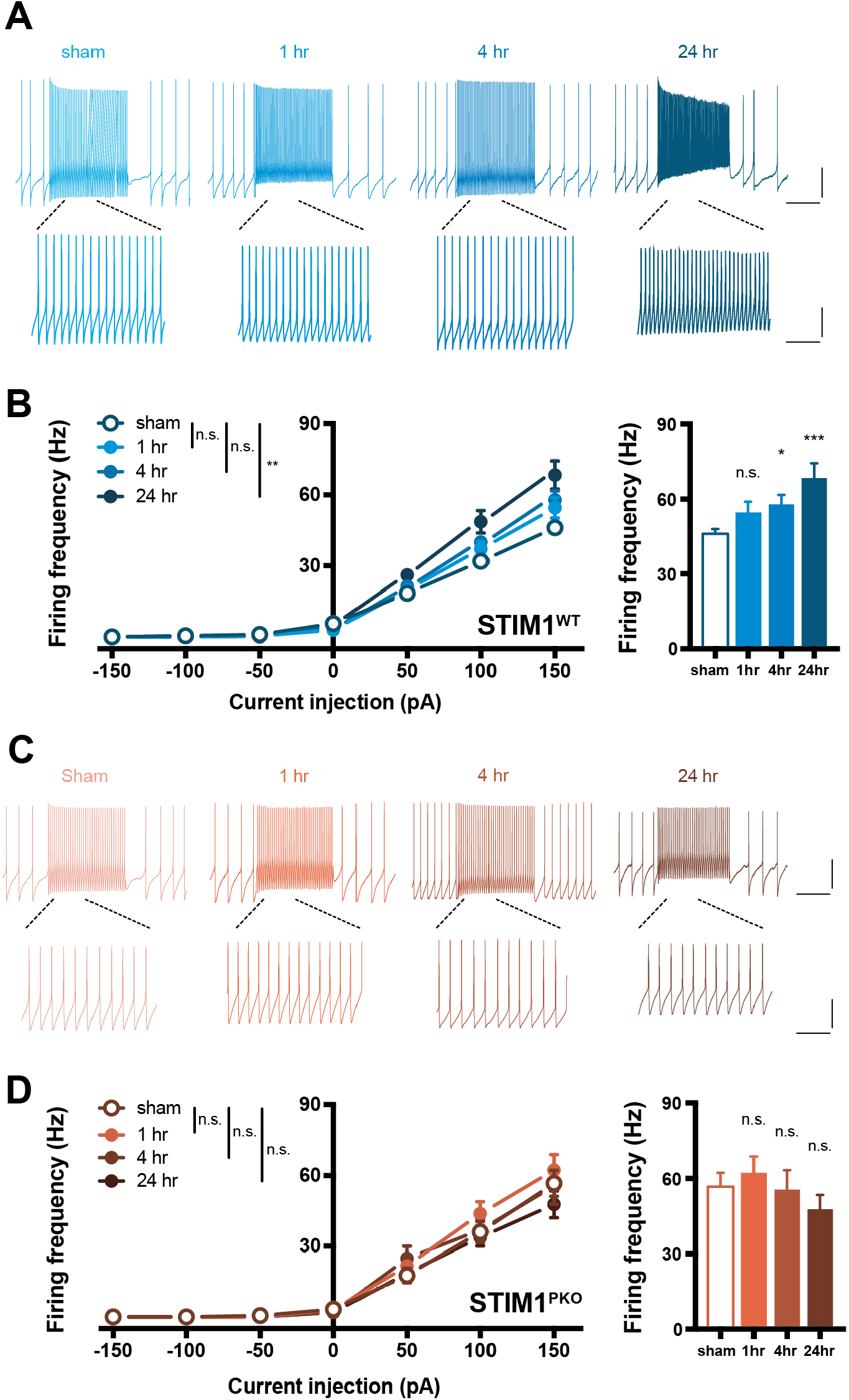
VOR gain-up learning induced potentiation of intrinsic excitability in VN neurons. (A) Representative traces from whole-cell recordings of wild-type group in each time point. Scale bars (upper), 20 mV (vertical) and 500 ms (horizontal). Scale bars (lower), 20 mV (vertical) and 125 ms (horizontal). (B) The excitability of VN neurons in wild-type littermates. VOR adaptation significantly potentiated gain responses of the VN neurons in response to square-wised current injection ranging from −150 pA to 150 pA for 1 s (sham vs 1hr, p=0.319; sham vs 4hr, p=0.072; sham vs 24hr, p=0.002, left; sham, n=23; 1hr, n=16; 4hr, n=38; 24hr, n=19). Excitability in 150 pA injection was significantly increased at mid-(4hr, p=0.032) and long-term (24hr, p < 0.001) after training. (C) Representative traces from whole-cell recordings of wild-type group in each time point. Scale bars (upper), 20 mV (vertical) and 500 ms (horizontal). Scale bars (lower), 20 mV (vertical) and 125 ms (horizontal). (D) Excitability of VN neurons in STIM1^PKO^. There was no alteration of excitability after learning (sham vs 1hr, p=0.422; sham vs 4hr, p=0.801; sham vs 24hr, p=0.493, left; sham, n=17; 1hr, n=25; 4hr, n=15; 24hr, n=15), and no significant changes in 150 pA injection as well (1hr, p=0.530.; 4hr, p=0.908; 24hr, p=0.371) Two-way repeated measure ANOVA was used for injected current-frequency graphs. One-way ANOVA with Fisher LSD test was used for bar graphs. Asterisks in each time points were calculated by comparing to sham groups. Error bars denote SEM. *p < 0.05 ***p < 0.001.

## Discussion

Here, we demonstrate a role of the intrinsic plasticity of cerebellar Purkinje cells in motor learning through which the VOR adaptive memory is transferred to the sub-cortical region for long-term memory storage. After gain-up training, the synaptic strength at PF-PC synapses is decreased and this learning-induced PF-PC LTD occurs concomitantly with a reduction of intrinsic excitability in PCs. Furthermore, VOR adaptation causes potentiation of the synaptic weight and intrinsic excitability in VN neurons, as well as the plasticity in PCs. Mice that were subjected to impaired memory consolidation, the STIM1^PKO^ group, show a normal learning curve and synaptic plasticity, whereas the PC intrinsic plasticity is declined within an hour. In addition to the unstable induction of PC intrinsic plasticity, there were no appropriate learning-induced alterations of synaptic transmission and excitability in the VN neurons despite both synaptic and intrinsic plasticity were capable to be induced. Our observations indicate that experience-dependent modulation of the neuronal excitability is required for long-term memory consolidation, in terms of the cerebellum-dependent motor learning.

Despite many implications suggesting that long-term storage of motor memory requires the memory transfer process from cerebellar cortex to nuclei (Boyden et al., 2004; Ito, 2013; Okamoto et al., 2011a; Okamoto, Shirao, Shutoh, Suzuki, & Nagao, 2011b; Shutoh et al., 2006), the detailed mechanisms of this memory transfer have yet to be elucidated. The data we presented here provides experimental evidence that VOR training results in the alteration of synaptic weight and excitability at multiple sites, the cerebellar cortex and the VN. Moreover, this study elucidates an unrevealed role of the intrinsic plasticity of cerebellar PCs in the VOR memory circuit by using the memory consolidation deficit mice model (Figure 1). Both learning-induced synaptic and intrinsic plasticity at the PC are observed in wild-type littermates (Figure 2 and 3). However, intrinsic plasticity is abolished within an hour of the learning task in the memory consolidation deficient mouse model (Figure 3), even though synaptic plasticity was normally induced (Figure 2). Furthermore, this impairment of the intrinsic plasticity of PCs is concomitant with the failure of VOR training-induced plasticity induction in the VN neurons (Figure 4), although the VN neurons in the STIM1^PKO^ mice are endowed with neural plasticity *in vitro* (Figure 5-figure supplement 1), implying that the learning-induced alteration of excitability in the PCs might serve as an instructive signal to induce the appropriate plasticity induction in VN neurons (Figure 6). These results support the previous expectations that PC activity can affect the synaptic and intrinsic plasticity induction in VN neurons (McElvain et al., 2010). Collectively, our results reconcile two long-standing hypotheses by providing experimental evidence for the induction of multiple forms of plasticity through VOR adaptation in both the cerebellar cortex and the brainstem.

**Figure 6.**
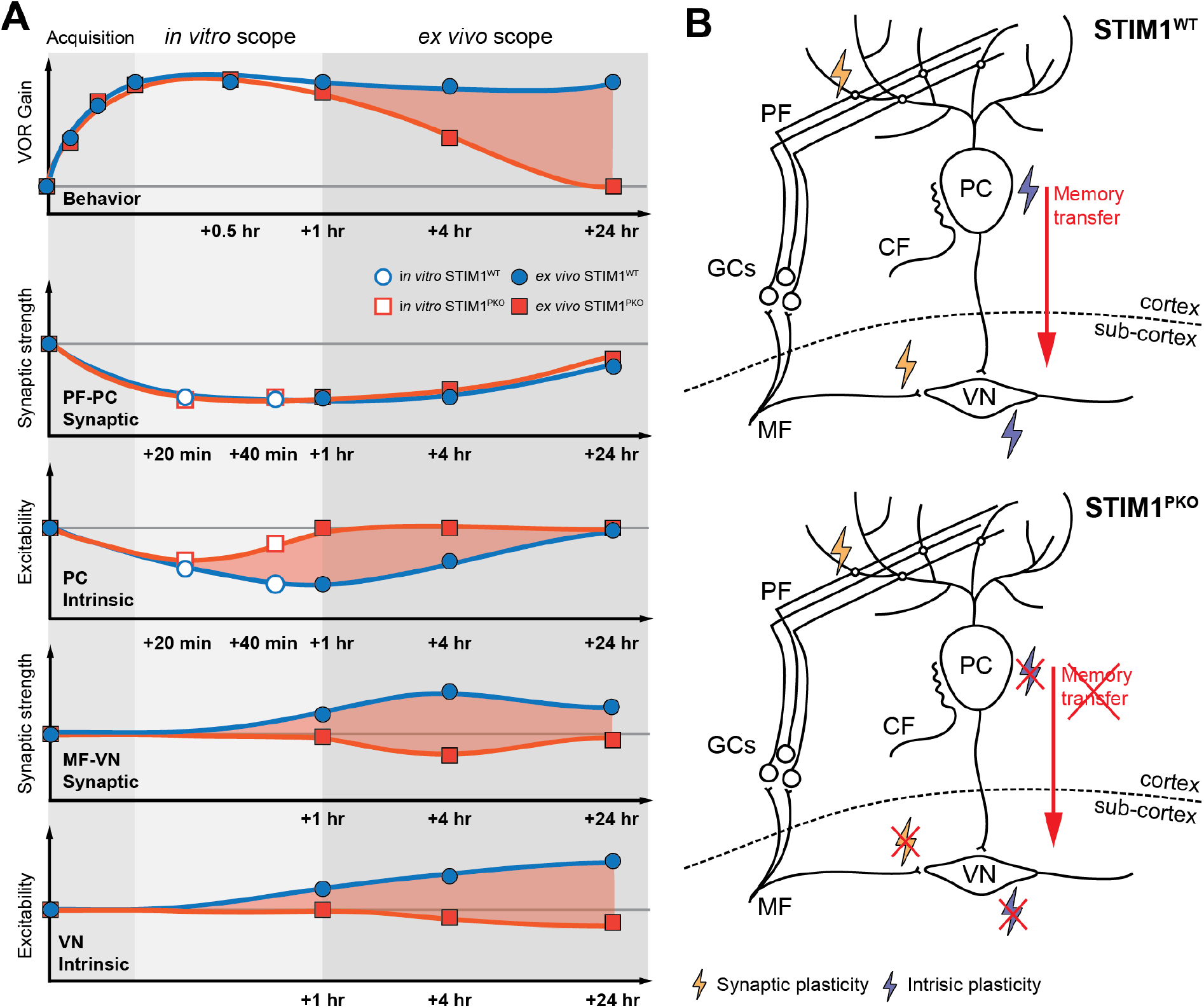
Schematic illustration for memory trace of VOR motor memory. (A) Summary of behavioral test and *in vitro*, *ex vivo* recording showing temporal order of the memory retention and neural plasticity in PCs and VN neurons. Differences between wild-type and STIM1^PKO^ mice are indicated as red shade of each plots). VOR memory retention level is maintained over a day whereas the motor memory is declined at mid-term period (+1hr to + 4hr) and long-term (+24hr) in STIM1^PKO^. Alterations of the neural activity corresponding to each period are presented below. There is no difference in PF-PC synaptic plasticity between wild-type and STIM1^PKO^ mice. However, learning-induced LTD-IE is abolished within an hour in STIM1^PKO^ and the difference in PC intrinsic plasticity between groups may lead to MF-VN synaptic plasticity and intrinsic plasticity of VN neurons (see the red shade). Furthermore, peak difference of each plot seems to move from 1 to 24 hours after learning, indicating that the plasticity in cerebellar PCs and VN neurons is connected in order. Thus, we speculate the PC intrinsic plasticity, which plays a role in memory transfer process, but not PF-PC synaptic plasticity underlies behavioural difference between wild-type littermates and STIM1^PKO^ (red shade in top). (B) Schematic illustration of neural circuit for VOR memory storage shown in wild-type (top) and STIM1^PKO^ mice (bottom). For successful memory acquisition and storage, four different types of neural plasticity are necessary (synaptic and intrinsic plasticity in the PCs and VN neurons). Especially, intrinsic plasticity of PC has important role in transfer acquired memory to sub-cortical area in this circuitry (top). When PC intrinsic plasticity is abolished, synaptic and intrinsic plasticity in VN neurons are impaired, thereby failure to transfer the acquired memory from the cortex to sub-cortex (bottom).

It has been assumed that motor memory is firstly formed in the cerebellar cortex and that neurons in the VN are involved in late phase adaption for VOR gain (Ito, 2013; Shutoh et al., 2006). This assumption implies that the temporal order between PC and VN plasticity has to be considered in memory processing. In our results, the VN plasticity is induced at a relatively later period than the plasticity in the PCs, and it indicates two major aspects. One is that PF-PC LTD contributes to memory acquisition, and the other is that the consequent induction of plasticity in VN neurons encodes long-term memory storage. Our data indicates that the impaired intrinsic plasticity of the cerebellar PCs would impair memory transfer and disrupt long-term memory storage. This supports the theory that the intrinsic plasticity of PCs connects two distinct brain regions and shapes the flow of information flow from the cerebellar cortex to the brainstem. The temporal order of plasticity at multiple sites may reflect the loci of memory storage. The *ex-vivo* recordings we presented here were executed at distinct time points: short-(∼1hr), mid-(∼4hrs) and long-term (∼24hrs) periods after learning. At the short-term period, the VOR adaptation curve and synaptic plasticity were not impaired in the memory consolidation deficient mouse model, although intrinsic plasticity was abolished (Figure 6). These results indicate that the memory acquisition may require synaptic plasticity in the cerebellar cortex, but not intrinsic plasticity. Rather, the aspects of the memory retention and deficiency of intrinsic plasticity in STIM1^PKO^ lead us to assume that the learning-induced alteration in PC excitability might be involved in the memory transfer process. Consistent with previous implications, our results suggest that the memory transfer occurs within 4 hours after learning (Kassardjian et al., 2005; Okamoto et al., 2011a; Shutoh et al., 2006). Synergies between synaptic and intrinsic plasticity may provide an instructive signal to convey the learned information into the brainstem, the VN, at the mid-term (∼4hrs) period. Interestingly, the synaptic plasticity in the VN neurons is observed slightly later than the intrinsic plasticity of the PC. Additionally, there is another slight delay in the VN intrinsic plasticity to reach peak (Figure 6). These results indicate that sequential flow of information from the cerebellar cortex to the sub-cortical region is responsible for memory processing. Taken together, we conclude that the acquired VOR memory might be located in the cerebellar cortex and the VN at the short- and long-term period, respectively, and a guiding instructive signal, driven by the intrinsic plasticity of the PCs may take part in the transfer of memory from the cortical area to the brainstem during the mid-term time period.

It is widely believed that the plasticity of neuronal excitability is involved in the cellular mechanism for memory storage (Daoudal & Debanne, 2003; Zhang & Linden, 2003). In particular, the intrinsic plasticity of cerebellar PCs shows features in the cerebellar memory circuits that are distinct from other types of neurons. In the neurons in the amygdala and hippocampal, learning-related neurons show higher excitability (Zhou et al., 2009), and the depolarization of the membrane potential of these cells enables the promotion of further synaptic plasticity (Ramakers & Storm, 2002; Watanabe, Hoffman, Migliore, & Johnston, 2002). Thus, these excitable neurons form a stable connection by strengthening the synaptic weight the given neural network, thereby consolidating the memory. In contrast, previous study suggested that the intrinsic plasticity of PCs occludes the subsequent induction of PF-PC synaptic plasticity (Belmeguenai et al., 2010). Hence, the plasticity of excitability may ensure that synaptic activity remains within a physiological limit by restricting further synaptic plasticity and adjusting the impact of PF activation on the output of PCs. In addition, our data show that there is no significant difference in the magnitude of synaptic plasticity at the PF-PC synapses between the wild-type littermate group and STIM1^PKO^ group, although the excitability is lower in the STIM1^PKO^ group than the wild-type group (Figure 3-figure supplement 1 and Figure 4-figure supplement 1A). Furthermore, the learning curve of VOR gain-up training was comparable between STIM1^PKO^ and wild-type littermates in spite of noticeable differences of basal excitability in the PCs among genotypes. Altogether, these results suggest that the basal membrane excitability in PCs is not thought to be correlated with the synaptic plasticity induction or the magnitude of synaptic plasticity. Considering the changes of neuronal activity in VN neurons over time, mid-term (∼4hrs) may critical period to induce the neural plasticity in VN. Interestingly, the PC intrinsic plasticity was found to be vanished within an hour after learning (Figure 3-figure supplement 1). We previously described that the concurrence of bidirectional synaptic and intrinsic plasticity may synergistically shape the cerebellar PC output (Shim et al., 2017). In this scenario, impairment of PC intrinsic plasticity shown in STIM1^PKO^ would break down the synergies between synaptic and intrinsic plasticity thereby deficit in plasticity in VN neurons. In fact, the sufficient cerebellar PC output was found to be exhibited when both synaptic and intrinsic plasticity in PCs occur (BioRxiv citation). Beside, individual activity of PC or the synergistic effect, the temporal synchrony of PCs output is considerable factor that contributing to information delivery from PC to DCN (Person & Raman, 2011). Accordingly, our data support the idea that activity-dependent alteration of synaptic weight and excitability in PCs play a role in providing an instructive signal contributing to plasticity in VN neurons. In conclusion, we suggest that learning-induced intrinsic plasticity may amplify the alteration of the synaptic transmission, resulting in the synergistic modulation of the net output of PCs in order to maximize information storage.

## Materials and Methods

### Animal model

We crossed homozygous PCP2-Cre line (B6.129-Tg(Pcp2-cre)2Mpin/J line, Jackson Laboratory) with the STIM1-floxed line (C57BL/6 background) to generate PC-specific STIM1 knockout line as our recent study (Ryu et al., 2017). Only male mice were used in all experiments, and procedures were approved by the Institutional Animal Care and Use Committee of Seoul National University College of Medicine.

### Surgery

All surgical procedure was similar to our previous paper (see (Ryu et al., 2017)). Surgery for head fixation has been done at 7- to 8- weeks-old. Zoletil (Zoletil 50, Virbac, 15mg/kg) + xylazine (Rompun, Bayer, 15mg/kg) mixture was applied to anesthetize mice through intraperitoneal injection. After surgery, 24 to 48 hours of recovery time were given to mice.

### Behavior test

The installation and most of procedure were similar to our previous paper (see (Ryu et al., 2017)). To control pupil dilatation, physostigmine salicylate solution (Eserine; Sigma Aldrich) was treated with brief isoflurane anesthetization, and at least 20mins were given to washout the side-effect of anesthetization. The concentration of eserine solution was constantly increased from 0.1%, 0.15% and 0.2% because of drug resistance. Two sessions of acclimation were performed. At first, the mouse was fixed onto a restrainer for 15 minutes and experienced light on and off, several brief visual and vestibular stimulation for checking surgical failure. Secondly, calibration step was included. We measured three basal ocular-motor responses, which are OKR, dVOR and lVOR. Visual stimulation via drum rotating was provided in sinusoidal rotation with ±5° for OKR. Vestibular stimulation was delivered through turn table rotation for dVOR and lVOR with same rotation amplitude as OKR. The only different between dVOR and lVOR was under light off and on, respectively. Each response was recorded at four different rotating frequencies 0.1, 0.25, 0.5, 1.0Hz. To increase 0.5Hz dVOR gain, associative visuo-vestibular stimulation was applied. Drum and table simultaneously rotated with ±5° of amplitude out of phase. The protocol contained three 10min training sessions and four check points (Figure 1A). Trained mice went back to its home cage which placed in complete dark condition until next check point, and after examine memory retention level, the mouse was sacrificed for *ex vivo* recording. Given stimulus and the response were fitted to sine curves. Gain was calculated by the ratio of the response amplitude to stimulus amplitude, and the phase differences between the two sine curves were determined as phase values. Custom built LabView (National Instrument) analysis tool was used for all these procedures. The level of memory consolidation was calculated by percent ratio of remained memory to learned memory.

### Slice preparation

Coronal cerebellar (flocculus) and brainstem slices of 270 - 320 µm were dissected by vibratome (Leica, VT1200) from behaviour tested 9 to 11 weeks old male mice in ice-cold NMDG cutting solution contained with the following (in mM): 93 NMDG, 93 HCl, 2.5 KCl, 1.2 NaH_2_PO_4_, 30 NaHCO_3_, 20 HEPES, 25 Glucose, 5 sodium ascorbate, 2 Thiourea, 3 Sodium pyruvate, 10 MgSO_4_·7H_2_O, 0.5 CaCl_2_·2H_2_O (pH 7.3). The brainstem slices containing VN were obtained from more rostral part in which the brainstem was attached to the cerebellum. The coronal plane of the cerebellar and brainstem slices were transferred into recovery chamber containing NMDG-cutting solution at 32 °C for 10 minutes, and then incubated in standard artificial cerebrospinal fluid (aCSF) contained with the following (in mM): 125 NaCl, 2.5 KCl, 1 MgCl_2_, 2 CaCl_2_, 1.25 NaH_2_PO_4_, 26 NaHCO_3_, 10 glucose at room temperature for an hour. NMDG-cutting solution and aCSF were oxygenated with 95% O_2_-5% CO_2_ (pH 7.4).

### Whole cell recording

Brain slices were put onto a submerged recording chamber on the stage of Olympus microscope (BX50WI, Japan) and perfused with standard aCSF. We used EPC9 amplifier with PatchMaster software (HEKA Elektronik) and multiclamp 700B amplifier with pClamp 10 (Molecular Device). Sampling frequency of 20 kHz and signals were filtered at 2 kHz (1 kHz filter for sEPSC). Inhibitory synaptic inputs were totally blocked by 100 µM picrotoxin (Sigma) in PC recording, and strychnine (1 µM) was added to block glycinergic input in VN recording. Patch pipettes (3-4 MΩ) were borosilicate glass and filled with internal solution containing the following: 9 KCl, 10 KOH, 120 K-gluconate, 3.48 MgCl_2_, 10 HEPES, 4 NaCl, 4 Na_2_ATP, 0.4 Na_3_GTP and 17.5 sucrose (pH 7.25) for testing *in vitro* recordings, *ex vivo* PC excitability and *ex vivo* VN recordings; 140 CsCl, 4 NaCl, 0.5 CaCl_2_, 10 HEPES, 2 MgATP and 5 EGTA (pH 7.3) for *ex vivo* sEPSC recording from PCs in the medial part of the flocculus. To evaluate the excitability of the cerebellar Purkinje cells, we compared gain responses of the neurons. We injected series of square-wised current steps ranging from +100 pA to +600 pA with increments of 100 pA for 500 ms. The membrane potential of the cerebellar Purkinje cells was held at −70 mV by injecting bias current in current clamp mode. To measure the excitability of VN neurons, we injected from −150 pA to +150 pA with increments of 50 pA for 1 s. There was no bias current in current clamp mode to hold membrane potential at these recordings. All patch clamp data, except for sEPSC recordings, were imported and analyzed by Igor Pro (Wave Metrics). The sEPSC data were analyzed using Mini Analysis (Synaptosoft). Other recording and analysis details including plasticity induction protocol were similar to our previous paper (see Shim et al., 2017)

#### Statistical analysis

As we described above, behavior data was analyzed by custom built LabView (National Instrument) tool, and electrophysiology data was analyzed by Igor Pro (Wave Matrics) and Mini Analysis (Synaptosoft). All statistical analysis was performed using Graphpad Prism 7 and Microsoft Excel. One-way ANOVA with Fisher’s LSD test, One-or Two-way repeated measure ANOVA with Sidak’s post-hoc test was used for several time groups analysis, and unpaired *t*-test was performed to compare wild-type and knockout group. All graphs are shown as mean±SEM, and asterisks *, ** and *** indicates p<0.05, p<0.01 and p < 0.001, respectively. *n* for each experiments are written in the figure legends.

## Acknowledgments

We thanks to Seungha Kim and Yong Gyu Kim for experimental support, and Misun Moon, Geehoon Chung for reviewing original draft. This study was supported by National Research Foundation of Korea (NRF) Grant funded by Korean government (MSIP) (2018R1A5A2025964, 2017M3C7A1029611 and 2016R1D1A1A02937282 to S.J.K; Global Ph.D fellowship programme, 2013H1A2A1034318 to D.C.J). H.G.S. received a scholarship from the BK21-plus education program provided by the National Research Foundation of Korea (5262-20170100)

## Author Contributions

Conceptualization, Investigation, Writing (Original Draft), D.C.J. and H.G.S.; Writing (Review & Editing), D.C.J., H.G.S. and S.J.K.; Funding Acquisition, D.C.J. and S.J.K.

---

**Figure 1-figure supplement 1.**
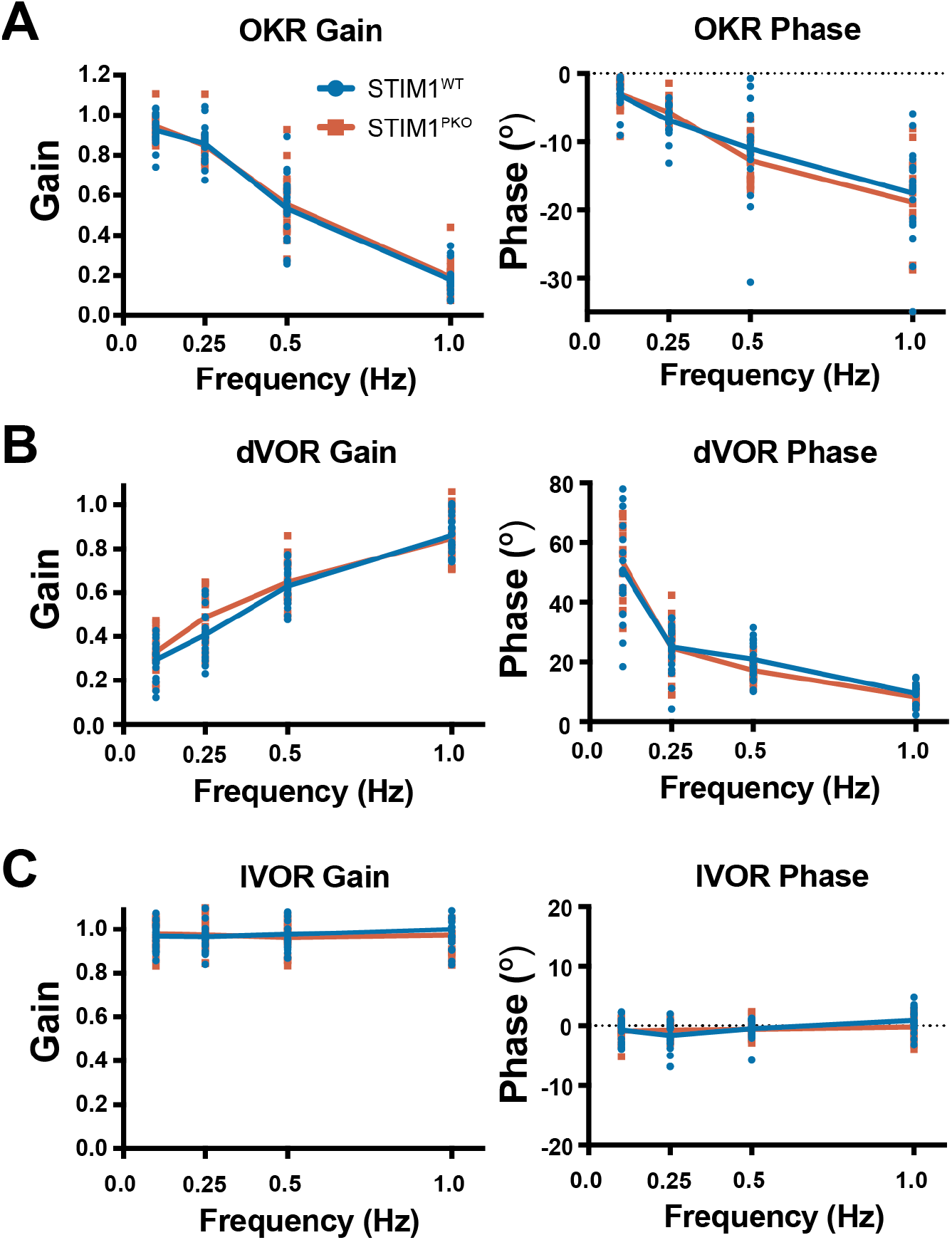
The basal ocular-motor performance of STIM1^PKO^ mice was not altered compared to wild-type littermates. (A) OKR responses in various frequency of drum rotating. OKR gain and phase values were not different between wild-type littermates (n=20) and STIM1^PKO^ (n=17). (B) VOR responses under dark (dVOR) in various frequency of drum rotating. dVOR gain and phase values were not different between wild-type littermates and STIM1^PKO^. (C) VOR responses under light (lVOR) in various frequency of drum rotating. lVOR gain and phase values were not different between wild-type littermates and STIM1^PKO^.

**Figure 2-figure supplement 1.**
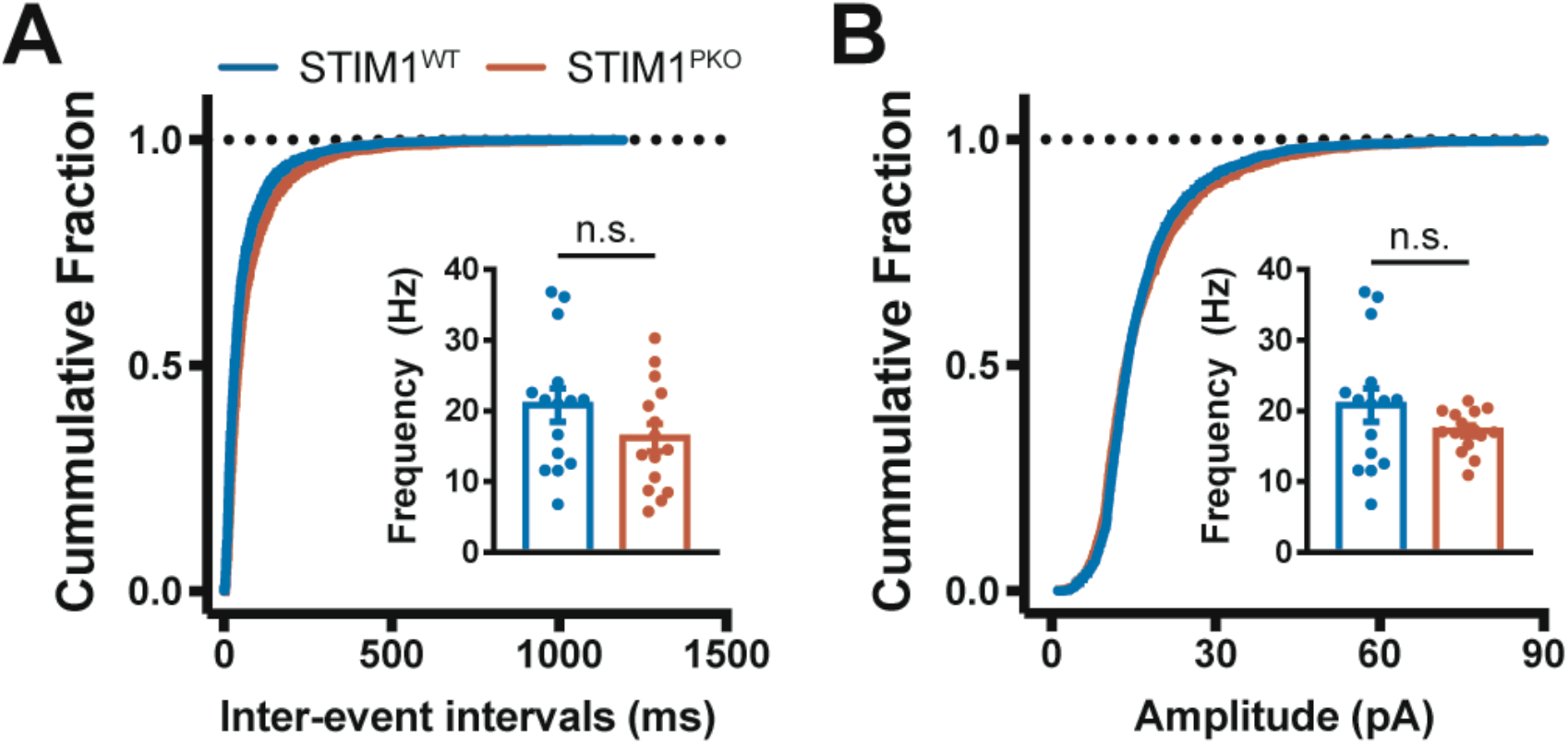
Synaptic transmission in STIM1^WT^ and STIM1^PKO^ was not significantly different. (A) Cumulative plots of IEI of sEPSC in wild-type (blue) and STIM1^PKO^ mice (red). The cumulative fraction of IEI and bar graph (inset) of sEPSC frequency indicated that frequency of sEPSC was not changed in STIM1^PKO^ compared to wild-type littermates (wild-type, n=15; STIM1^PKO^, n=15, p=0.145). (B) Cumulative plots of amplitude of sEPSC. The cumulative fraction of amplitude and bar graph (inset) of sEPSC frequency indicated that amplitude of sEPSC was not changed in STIM1^PKO^ compared to wild-type littermates (p=0.587). Unpaired *t*-test was used for bar graphs. Error bar denotes SEM.

**Figure 3-figure supplement 1.**
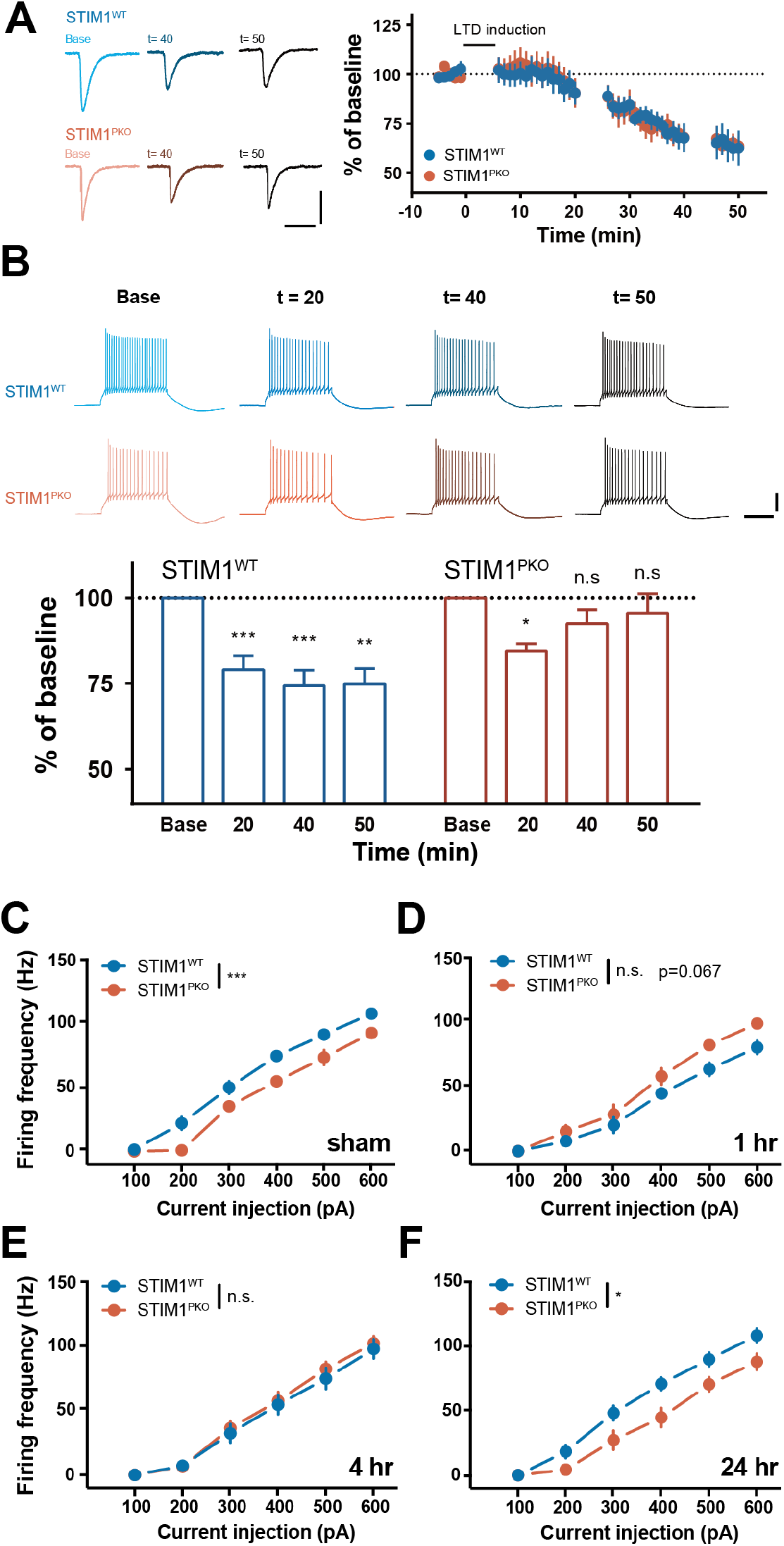
Intrinsic plasticity was impaired in STIM1^PKO^. (A) Plots of the normalized EPSC change after application of LTD induction protocol in wild-type littermates (n=8, n=4 after 40 min, blue) and STIM1^PKO^ mice (n=7, n=6 after 40 min, red). Synaptic plasticity was normally induced in STIM1^PKO^ mice. Scale bar, 200 pA (vertical) and 50 ms (horizontal) (B) Bar graphs showed the comparison of excitability changes after LTD induction. In STIM1^PKO^ mice, down-regulation of excitability in PCs was shown 20 min after induction (n=7, p=0.010), however, fully recovered 40 min and 50 min after induction (40 min, n=7, p=0.331; 50 min, n=6, p=0.748). On the contrary, the intrinsic plasticity was induced and slightly further developed in time from wild-type littermates (20 min, n=8, p < 0.001; 40 min, n=8, p < 0.001; 50 min, n=4, p=0.001). Scale bars (upper), 20 mV (vertical) and 200 ms (horizontal) (C-F) Comparing PC excitability of wild-type littermates (blue) and STIM1^PKO^ mice (red) in each time points. (C) STIM1^PKO^ group (n=17) showed significantly lower excitability than wild-type littermates (n=20, p < 0.001). (D) While the excitability of STIM1^PKO^ (n=13) was unchanged, excitability of wild-type littermates (n=11) was plunged (p=0.067). (E) At 4 hours after learning, the altered excitability of wild-type littermates (n=16) partly restored, and overlapped to excitability of STIM1^PKO^ (n=17), which was still unchanged (p=0.644). (F) As the excitability of wild-type littermates (n=20) was fully recovered, significant difference from STIM1^PKO^ group (n=13) was also restored (p=0.022). One-way repeated measure ANOVA with post hoc Sidak correction was used for panel B. Two-way repeated measure ANOVA was used for panel C to F. Error bars denote SEM. *p < 0.05, **p < 0.01, ***p < 0.001.

**Figure 4-figure supplement 1.**
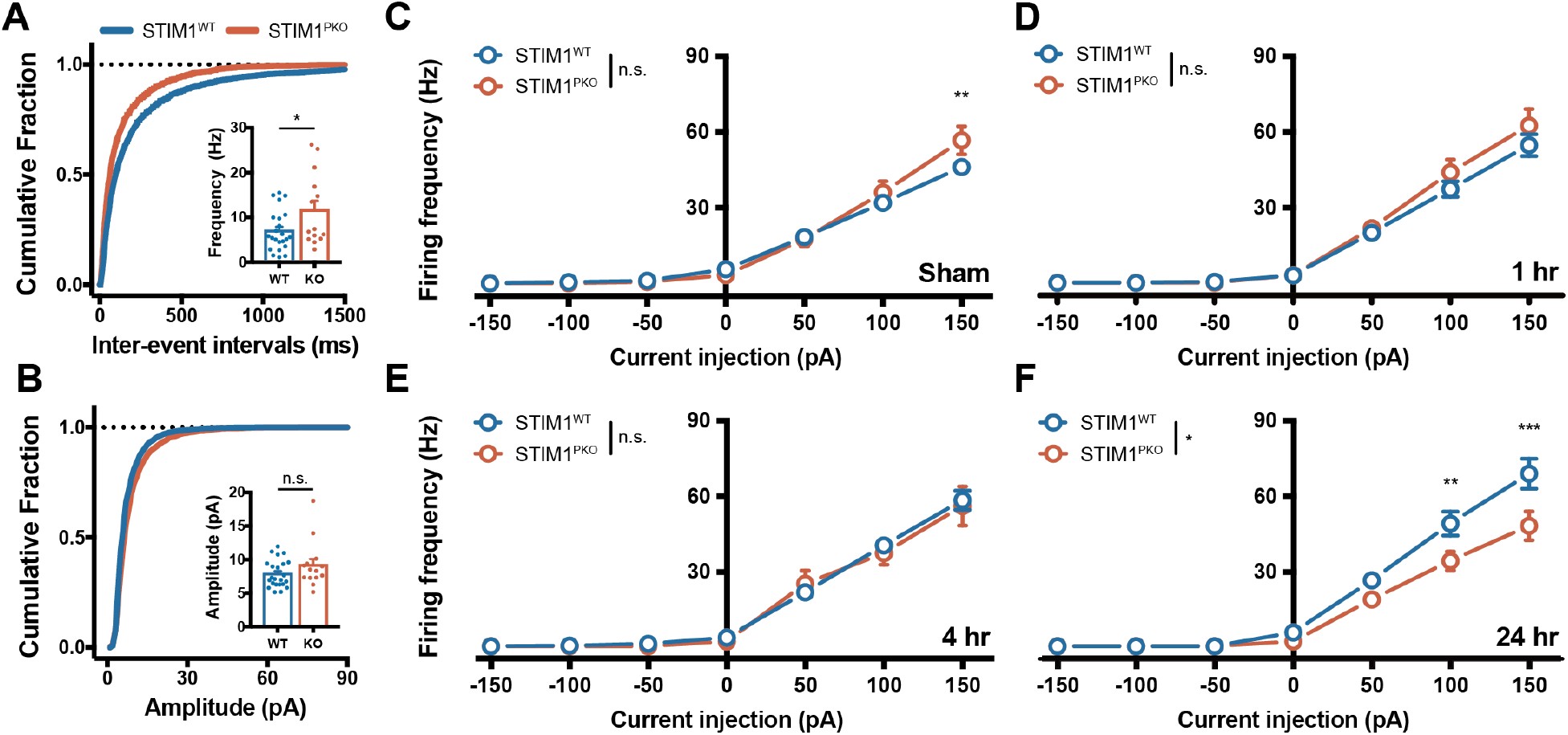
Basal synaptic transmission and excitability in VN neurons before VOR gain up training. (A) Frequency of synaptic transmission in VN neurons from wild-type littermates (n=23, blue) and STIM1^PKO^ mice (n=14, red). The cumulative fraction of IEI and bar graph (inset) of sEPSC frequency indicated that frequency of sEPSC was higher in STIM1^PKO^ compared to wild-type littermates (p=0.032). (B) Amplitude of sEPSC in VN neurons from wild-type littermates (blue) and STIM1^PKO^ mice (red). The cumulative fraction of amplitude and bar graph (inset) of sEPSC frequency indicated that amplitude of sEPSC was not changed in STIM1^PKO^ compared to wild-type littermates (p=0.161). (C-F) Excitability in VN neurons from STIM1^WT^ (blue) and STIM1^PKO^ (red) mice in each time points. Square-wised somatic current steps were injected from membrane potential with various ranges from −150 pA to 150 pA with increment of 50 pA for 1 s. (C) Overall, the gain responses of VN neurons from STIM1^PKO^ (n=16) were not significantly different from wild-type littermates (n=23, p=0.423), but at 150 pA injection, STIM1^PKO^ showed higher excitability than wild-type littermates (p=0.003). (D) At the short-term period, the excitability of VN neurons of the wild-type (n=16) has slightly increased, while the excitability of STIM1^PKO^ (n=25) was unchanged. Statistical difference has disappeared (p=0.428). (E) At 4 hours after learning, the excitability of VN neurons of the wild-type (n=38) increased more, and that of STIM1^PKO^ (n=15) was unchanged again. IO curve of both groups are overlapped (p=0.759). (F) As the excitability of wild-type littermates (n=19) became much higher at the long-term period, the wild-type group showed significant higher frequency than STIM1^PKO^ group (n=15) (p=0.016). At 100 pA and 150 pA injection, statistical significance was 0.005 (100 pA) and <0.001 (150 pA) by post-hoc Sidak test. Unpaired *t*-test was used for bar graphs in panel A and B. Two-way repeated measure ANOVA with Sidak test was used for panel C-F. Error bars denote SEM. *p < 0.05, **p < 0.01, ***p < 0.001.

**Figure 5-figure supplement 1.**
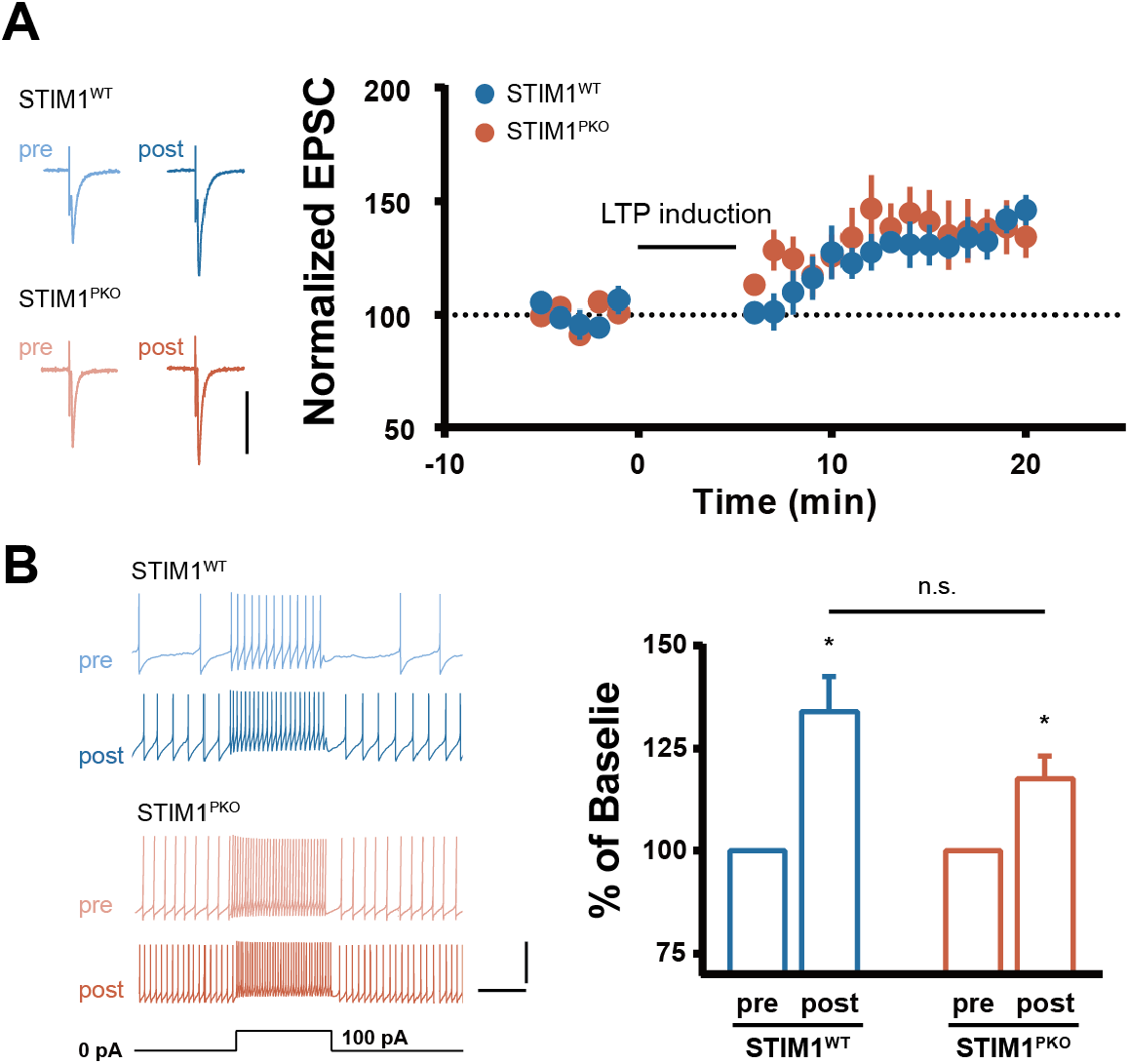
Both synaptic and intrinsic plasticity was able to be induced in both genotypes. (A) Synaptic LTP of VN neuron. Both wild-type (n=6) and STIM1^PKO^ (n=7) groups showed intact synaptic plasticity through *in vitro* induction protocol. (B) Intrinsic plasticity of VN neuron was intact in both genotypes. Through LTP induction protocol, the excitability of VN neurons was significantly increased in both genotypes (wild-type, n=6, p=0.031; STIM1^PKO^, n=6, p=0.031). There was no significant difference between post-induction groups of both genotypes (p=0.386) Mann-Whitney U test was used for panel B. Error bars denote SEM. *p < 0.05

